# Adult mice lacking VIP SCN neurons retain circadian locomotor behavior but exhibit dampened daily glucocorticoid rhythms

**DOI:** 10.1101/2020.01.07.897140

**Authors:** Cristina Mazuski, Samantha P. Chen, Erik D. Herzog

**Affiliations:** Department of Biology, Washington University, St. Louis, Missouri 63130-4899; Sainsbury Wellcome Centre, 25 Howland St, London W1T 4JG

**Author notes:** Conflict of Interest: The authors declare no competing financial interests.

## Abstract

The suprachiasmatic nucleus (SCN) drives circadian rhythms in locomotion through coupled, single-cell oscillations. Global genetic deletion of the neuropeptide, *Vip* or its receptor *Vipr2*, results in profound deficits in daily synchrony among SCN cells and daily rhythms in locomotor behavior and glucocorticoid secretion. To test whether this phenotype depends on VIP neurons in the SCN, we ablated VIP SCN neurons *in* vivo in adult mice through Caspase3-mediated induction of the apoptotic pathway in cre-expressing VIP neurons. We found that ablation of VIP SCN neurons in adult mice caused a phenotype distinct from *Vip-* and *Vipr2-* null mice. Mice lacking VIP neurons retained rhythmic locomotor activity with a shortened circadian period, more variable onsets and decreased duration of daily activity. Circadian hormonal outputs, specifically corticosterone rhythms were severely dampened. In contrast, deletion of neonatal SCN VIP neurons dramatically reduced circadian gene expression in the cultured SCN, mimicking the effects of global deletion of *Vip* or *Vipr2*. These results suggest that SCN VIP neurons play a role in lengthening circadian period and stimulating the daily surge in glucocorticoids in adults and in synchronizing and sustaining daily rhythms among cells in the developing SCN.

**Significance Statement:** The importance of the neuropeptide, VIP, for circadian rhythms has been described in mice lacking the gene for *Vip* or its receptor, *Vipr2*. This study found that ablation of VIP neurons only in the adult SCN reproduced the loss of circadian rhythms in glucocorticoids, but not the loss of circadian locomotor behavior, seen with global loss of VIP signaling. We conclude that VIP SCN neurons play two roles: one in adulthood lengthening circadian period and regulating circadian outputs, and one in development coordinating synchrony among circadian cells.

## Introduction

Located in the ventral hypothalamus, the suprachiasmatic nucleus (SCN) is the dominant pacemaker that aligns daily physiological and behavioral rhythms to the local light-dark schedule (Coomans, Ramkisoensing, & Meijer, 2015; Hastings, Maywood, & Brancaccio, 2018; Herzog, 2007; Welsh, Takahashi, & Kay, 2010). Individual SCN neurons express cell autonomous oscillations driven through a negative transcription-translation feedback loop involving the ‘core clock genes’, which include Bmal1, Clock, Period 1 and 2, and Cryptochrome 1 and 2 (Takahashi, 2017; Webb, Angelo, Huettner, & Herzog, 2009; Welsh, Logothetis, Meister, & Reppert, 1995). Intercellular neurotransmission is necessary to couple single-cell oscillators and generate high amplitude, precise, circuit-wide rhythms capable of driving behavioral circadian rhythms (Herzog, Aton, Numano, Sakaki, & Tei, 2004; Tokuda, Ono, Honma, Honma, & Herzel, 2018; Welsh et al., 1995; Yamaguchi et al., 2003).

Vasoactive intestinal polypeptide is sparsely expressed throughout the mammalian brain in the cortex, retina, superior colliculus and SCN. Released by 10% of SCN neurons, vasoactive intestinal polypeptide (VIP) is necessary for the synchronization of single-cell circadian rhythms (Abrahamson & Moore, 2001; Aton, Colwell, Harmar, Waschek, & Herzog, 2005). Mice lacking *Vip* or its receptor *Vipr2* show an advanced phase angle of entrainment in a light cycle and approximately 60% lose daily locomotor rhythms or exhibit multiple periodicities in constant darkness (Aton et al., 2005; Colwell et al., 2003; Cutler et al., 2003; Harmar AJ et al., 2002). *Vip-*null mice also lose rhythmicity in hormonal circadian outputs, for example the daily rhythm in corticosterone (Loh, Abad, Colwell, & Waschek, 2008). *In* vitro, SCN slices lacking VIP signaling (either through global genetic deletion of *Vip* or *Vipr2* or pharmacological blockade) show decreases in the number of rhythmic SCN neurons and the loss of intercellular synchrony (Aton et al., 2005; T. M. Brown, Colwell, Waschek, & Piggins, 2007; Maywood et al., 2006). These results led to the conclusion that VIP signaling is required to synchronize circadian cells and amplify daily rhythms in physiology and behavior. The relative roles of VIP neurons in the SCN, neocortex and other brain areas in the development and sustenance of circadian rhythms have not, however, been fully clarified.

Furthermore, mice that lack *Vip* or *Vipr2* display phenotypic heterogeneity in their circadian profile (only 60% of mice show behavioral arrhythmicity). This suggests that other non-VIP signaling pathways compensate for the loss of VIP signaling (Timothy M. Brown, Hughes, & Piggins, 2005; Maywood, Chesham, O’Brien, & Hastings, 2011). In Drosophila, the genetic knockout of pigment-dispersing factor (PDF, the functional homologue of VIP) yields a similarly disrupted circadian phenotype to the ablation of PDF-expressing neurons (Renn, Park, Rosbash, Hall, & Taghert, 1999). But cell ablation and genetic knockout need not produce identical phenotypes. For example, deletion of the Opn4 (melanopsin) gene has mild effects on circadian entrainment compared to the complete loss of photic entrainment following ablation of Opn4-expressing intrinsically photosensitive retinal ganglion cells (Güler et al., 2008; Hattar, Liao, Takao, Berson, & Yau, 2002). Therefore, we selectively ablated VIP SCN neurons *in vivo* and *in vitro* to test for their necessity in development and adulthood for circadian rhythms in locomotor activity, glucocorticoid release and gene expression.

Surprisingly, instead of a more severe phenotype, we found that ablation of VIP neurons *in* vivo in the adult mouse SCN yielded a mild circadian phenotype, primarily characterized by a shortened period during constant darkness, increased activity onset variability and shortened duration of activity. Despite this mild circadian phenotype, we found pronounced deficits in the circadian corticosterone rhythm. Interestingly, when VIP SCN neurons were ablated *in vitro* we found the hypothesized decreases in rhythmicity and amplitude. Our results suggest the intriguing possibility that in the intact adult brain, VIP neurons are not the sole conduit for synchronizing single-cell rhythms, but rather play a role in period lengthening and hormonal outputs.

## Materials and Methods

### Animals

All mice were housed in a 12h:12h light:dark cycle in the temperature- and humidity-controlled Danforth Animal Facility at Washington University in St. Louis with ad lib access to food and water. Combinations of the following genotypes were used in all experiments: VIP-IRES-Cre knock-in mice (VIP^tm1(creZjh)^, Jackson Laboratories, RRID: IMSR_Jax:010908), *Per2::Luciferase* knock-in mice (founders generously provided by Dr. Joseph Takahashi, UTSW), tdTomato reporter mice (B6.Cg-Gt(ROSA)26Sor^tm9(CAG-tdTomato)Hze^/J, Jackson Laboratories, #007909) and C57Bl/6JN mice. All procedures were approved by the Animal Care and Use Committee of Washington University and adhered to National Institutes of Health guidelines.

### In Vivo Deletion of VIP Neurons

pAAV-flex-taCasp3-TEVp plasmid was obtained from Addgene (gift from Nirao Shah & Jim Wells, Addgene plasmid #45580, http://n2t.net/addgene:45580; RRID:Addgene_45580) and packaged into a high titer AAV8 virus by the Washington University Viral Core. To ablate VIP SCN neurons *in vivo*, 6-10 week-old VIP-IRES-Cre heterozygous mice were injected with AAV8-Casp3-TEVP virus targeted to the SCN. Briefly, anesthetized mice (2% isofluorane) were placed into a stereotactic device and the virus (0.5 μL/side) was bilaterally injected into the SCN at +0.4mm anterior, ± 0.15 mm lateral and 5.6mm ventral to Bregma. Age-matched control mice were either VIP-IRES-Cre heterozygotes receiving artificial cerebrospinal fluid (ACSF containing in mM: NaCl, 125; KCl, 2.5; NaH_2_PO_4_, 1.25; NaHCO_3_, 25; CaCl_2_, 2; MgCl_2_, 1; dextrose, 25 ~310 mOsmol/1) injections or mice lacking Cre expression (C57Bl6/JN or PER2::Luc/+) receiving AAV8-Caspase3-TEVP injections. Mice received analgesic treatment during recovery from surgery.

### Locomotor Activity

Following recovery, mice were placed in custom-built cages to monitor running wheel activity (Clocklab, Actimetrics, Evanston, IL) in 12h:12h light:dark for at least 7 days and in constant darkness for at least 14 days. Experiments were run on 4 separate cohorts of mice. Locomotor activity was analyzed from at least one week of data using Clocklab software (Actimetrics) and custom-written Python scripts. Circadian period, amplitude and rhythmicity were calculated using chi-squared periodogram. A 8-h box filter was used to identify daily activity onsets and offsets to calculate cycle-to-cycle onset variability and the duration of daily activity. Custom python scripts were used to quantify total daily activity counts. All data were analyzed blinded to treatment and genotype.

### Circadian Corticosterone Analysis

Following locomotor activity recordings, a subset of the mice were transferred to custom-built cages for fecal matter collection. These cages consisted of a wire stage where mice could comfortably sit with easy access to ad lib food and water. The majority of feces and urine fell through the wire stage onto a piece of filter paper at the bottom of the cage. A slot was cut out of the bottom front of the cage that allowed an investigator to easily collect and replace the filter paper. We estimate 85-100% of feces per time point were collected. After mice habituated to the cages in constant darkness for two days, investigators collected fecal samples in either 3- or 4-h time bins for 48-72h. Samples were immediately transferred to −80°C. For steroid extraction, samples were baked at 50°C for 3-4 h until completely dry before being individually ground into a powder using a mortar and pestle. Total weight per time point was calculated and steroids were extracted from 25mg of powdered fecal matter using 80% methanol. After agitation, the supernatant was transferred to new tubes and methanol was evaporated inside a fume hood. The pellets were suspended in 500 ul of ELISA buffer and diluted to a concentration of 1:2500 in ELISA buffer. Samples were processed in duplicate for corticosterone concentration using the ELISA kit instructions (Cayman Chemicals, Corticosterone EIA). Final concentrations (in ng per mg of feces) were determined based on the standard curve, dilution factor and total fecal weight per time point. There were no measurable differences in fecal weight between VIPN ablated and control mice. Corticosterone collection and measurement was completed on 3 separate cohorts of mice.

### Immunohistochemistry

Following locomotor and corticosterone assessment, mice were re-entrained to 12h:12h light:dark cycle and then perfused during late day (Zeitgeber Time, ZT 7-10). Briefly, mice were anesthetized with 1.25% Avertin (2,2,2-tribromoethanol and tert-amyl alcohol in 0.9% NaCl; 0.025 ml/g body weight) and transcardially perfused with phosphate-buffered saline (PBS) and 4% paraformaldehyde (PFA). The brain was dissected and transferred to 30% sucrose the following day. Frozen coronal sections were cut at 40μm on a cryostat (CM1850, Leica) and placed in 3 separate wells. For VIP and AVP immunofluorescence, free-floating sections were washed for 1 h in PBS, then blocked in PBSGT (5% normal goat serum, 0.3% Triton and PBS) for 1h. Sections were incubated overnight in primary antibodies diluted in PBSGT (rabbit anti-VIP 1:2000, Immunostar and mouse anti-AVP PS41 1:100, generous gift of Dr. Hal Gainer, NIH). Slices were washed again and incubated for 2 h at room temperature with secondary antibodies (anti-rabbit 488 and anti-mouse 564, 1:500). Sections were briefly stained with DAPI, washed again in PBS, mounted and cover-slipped with DABCO (1,4-Diazobicycol[2,2,2]-octane) mounting medium. All sections from a specific cohort were simultaneously processed. Sections were imaged on an epifluorescent microscope (TE2000 inverted, Nikon). The SCN was identified using the DAPI staining resulting in 2-5 SCN sections per animal. Images of VIP and AVP immunofluorescence were acquired using the same exposure and gain for every mouse in each round, allowing us to directly compare staining intensities. An investigator, blinded to the genotype and treatment of the mouse, drew boundaries around the SCN using ImageJ software and measured the mean intensity of VIP and AVP staining within those boundaries for each SCN section. Background staining for VIP and AVP was calculated as the mean intensity for each image from an area slightly lateral to the SCN that had no visible AVP or VIP labelling. To calculate relative staining intensity of VIP or AVP, the respective background staining was subtracted from the SCN VIP or AVP mean staining intensity. Values reported are the highest relative staining intensities for individual mouse. A subset (n = 7) of mice that received *in* vivo Credependent caspase3 injections expressed tdTOMATO in VIP neurons. Instead of immunohistochemistry, these mice were sectioned and imaged using fluorescence microscopy. Of these 7 mice, 1 had no tdTOMATO expression indicating ablation of VIP neurons (VIPN ablated), 3 retained VIP expression (VIPN present) and 3 lacked Vip-driven Cre expression (C57Bl/6JN controls). These 7 mice were included in Figure 1, but excluded from Figure 3.

**Figure 1.**
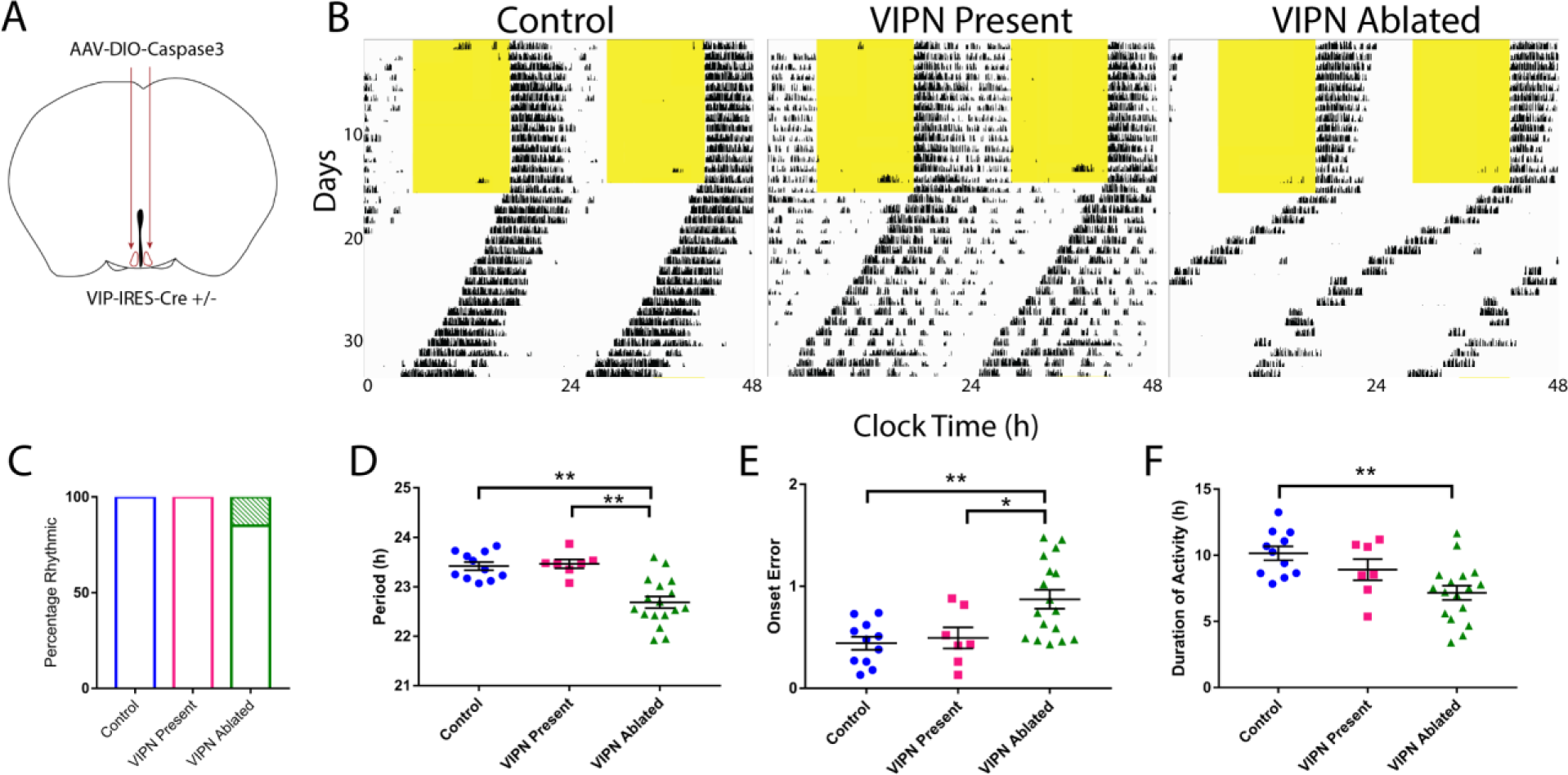
Ablation of SCN VIP Neurons in adult mice shortens the duration and period of daily locomotor activity. **A)** Schematic showing injection of Cre-dependent Caspase3 virus bilaterally into the SCN of adult VIP-Cre heterozygous mice. **B**) Representative actograms of locomotor activity in LD and DD for the following three groups: Control, VIPN present and VIPN ablated. **C**) Ablation of VIP SCN neurons had little effect on the percentage of circadian mice (3/20 VIPN Ablated mice were arrhythmic compared to 0/11 control and 0/7 VIPN present mice). (**D**) Ablation of VIP SCN neurons shortened circadian period, (**E**) increased onset variability, (**F**) and reduced the duration of daily activity. Error bars show mean ± SEM.

### In Vitro Deletion of VIP Neurons

The SCN from triple transgenic mice heterozygous for VIP-IRES-Cre, Rosa-tdtomato, and PER2::Luc were used for these experiments. Briefly, between postnatal day 6-7 (P6-P7), these mice were decapitated, and brains were rapidly removed. The bilateral SCN was dissected from 300-μm thick coronal brain slices, imaged to verify tdtomato fluorescence, and cultured on Millicell-CM inserts (Millipore, Billerica, MA) in pre-warmed culture medium (DMEM supplemented with 10mM HEPES and 10uM beetle luciferin, Promega, Madison, WI). The following day SCN were infected with 1-2μl of either AAV8-Caspase-TEVP or AAV9-DIO-eYFP viruses. The virus was applied directly on top of the slice. Each SCN explant in its sealed 35-mm Petri dish (BD Biosciences, San Jose, CA) was placed under a photomultiplier tube (HC135-11, Hamamatsu Corp.) in a light-tight incubator kept at 36°C. Bioluminescence counts were integrated every 10 minutes for up to a month using custom software. Each slice was imaged on an epifluorescent microscope (TE2000 inverted, Nikon) once per week to visualize the changes in number of VIP neurons and media was replaced every two weeks. All data were analyzed blinded to genotype and treatment. Circadian period was characterized using a cosine fit on detrended data (Chronostar software, generous gift of Dr. Achim Kramer, Charité University). Amplitude was calculated using a custom-written python script which measured the daily peak-to-trough difference from the raw data. Epifluorescent images for tdtomato-positive neurons were quantified using ImageJ’s cell counter plug in.

### Experimental Design and Statistical Analyses

All statistical tests were performed using Prism version 7.00 for Windows 7 (GraphPad Software). For circadian locomotor experiments, one-way ANOVAs were calculated to compare 3 groups (Control, VIPN present, and VIPN ablated). When comparing locomotor behavior between 4 groups (Control, VIPN present, VIPN ablated and *Vip* or *Vipr2* KO), we used either a one-way ANOVA or Kruskal Wallis test (e.g. when standard deviations were significantly different between the groups in comparing rhythmic amplitudes in Figure 3 and as noted in Table1). When comparing corticosterone measurements, VIP/AVP staining intensity, and bioluminescence, we used an un-paired Student’s t-test. Welch’s correction was applied when standard deviations were significantly different (corticosterone maximum and amplitude, AVP staining intensity, *in vitro* VIP neuron count, mean bioluminescence, correlation coefficient). We calculated the Pearson correlation to test the relationship between locomotor phenotype and VIP staining intensity.

## Results

### Ablation of VIP neurons in adult mice alters circadian locomotor activity

To test whether VIP SCN neurons are necessary for rhythmic locomotor behavior, we targeted them for ablation in adult mice and monitored running wheel activity. To selectively ablate VIP SCN neurons, we bilaterally injected an adeno-associated virus expressing Cre-dependent Caspase3, AAV8-Casp3-TEVP (Yang et al., 2013) into the SCN of VIP-IRES-Cre heterozygous mice (Figure 1A). Expression of the Caspase3 construct cell-autonomously triggered the apoptotic pathway resulting in selective ablation of VIP SCN neurons. At the conclusion of behavioral experiments, mice were sacrificed to confirm ablation of VIP SCN neurons. Hereafter, experimental mice with confirmed ablation of VIP neurons are referred to as “VIPN ablated” (n = 20). Virus-treated mice that retained wild-type levels of VIP expression are referred to as “VIPN present” (n = 7). Control mice were either VIP-IRES-Cre mice injected with ACSF (n = 5) or non-Cre expressing mice injected with AAV8-Casp3-TEVP (n=6).

Starting at least one week following surgery, we recorded wheel running behavior in entrained conditions (12h:12h light:dark, LD, cycle) for 1-2 weeks and then in constant darkness (DD) for 2-3 weeks (Figure 1B and Figure 2). Surprisingly, VIPN ablated mice showed normal entrainment in the light cycle and retained circadian rhythms in DD (Figures 1C, 2 and 3). While still rhythmic, mice lacking SCN VIP neurons had notably distinct phenotypes compared to controls in constant conditions, running for approximately 3 h less per day (7.17±0.54 h compared to 10.16±0.52 h, F(2,32) = 7.24, p = 0.0025), with a 0.7 h-shorter circadian period (22.69±0.12h compared to 23.42±0.08h, F(2,32) = 16.59, p <0.0001) and doubled variability in day-to-day activity onset (0.87±0.09h compared to 0.44±0.06h, F(2, 32) = 7.374, p = 0.0023, Table 1 and Figures 1D-F). These results were in striking contrast to *Vip-* and *Vipr2*-null mice which did not show these traits, but had reduced circadian amplitude (3409±1166 compared to 7304±3363 in VIPN ablated mice, H(2) = 16.58, p = 0.0003) and 50% (n=4 of 8), rather than 15% (n=3 of 20), were scored as arrhythmic in constant darkness (Figure 3A-B). Loss of VIP signaling from birth, thus, greatly impaired circadian rhythm generation which was intact following loss of VIP neurons in adulthood.

**Figure 2.**
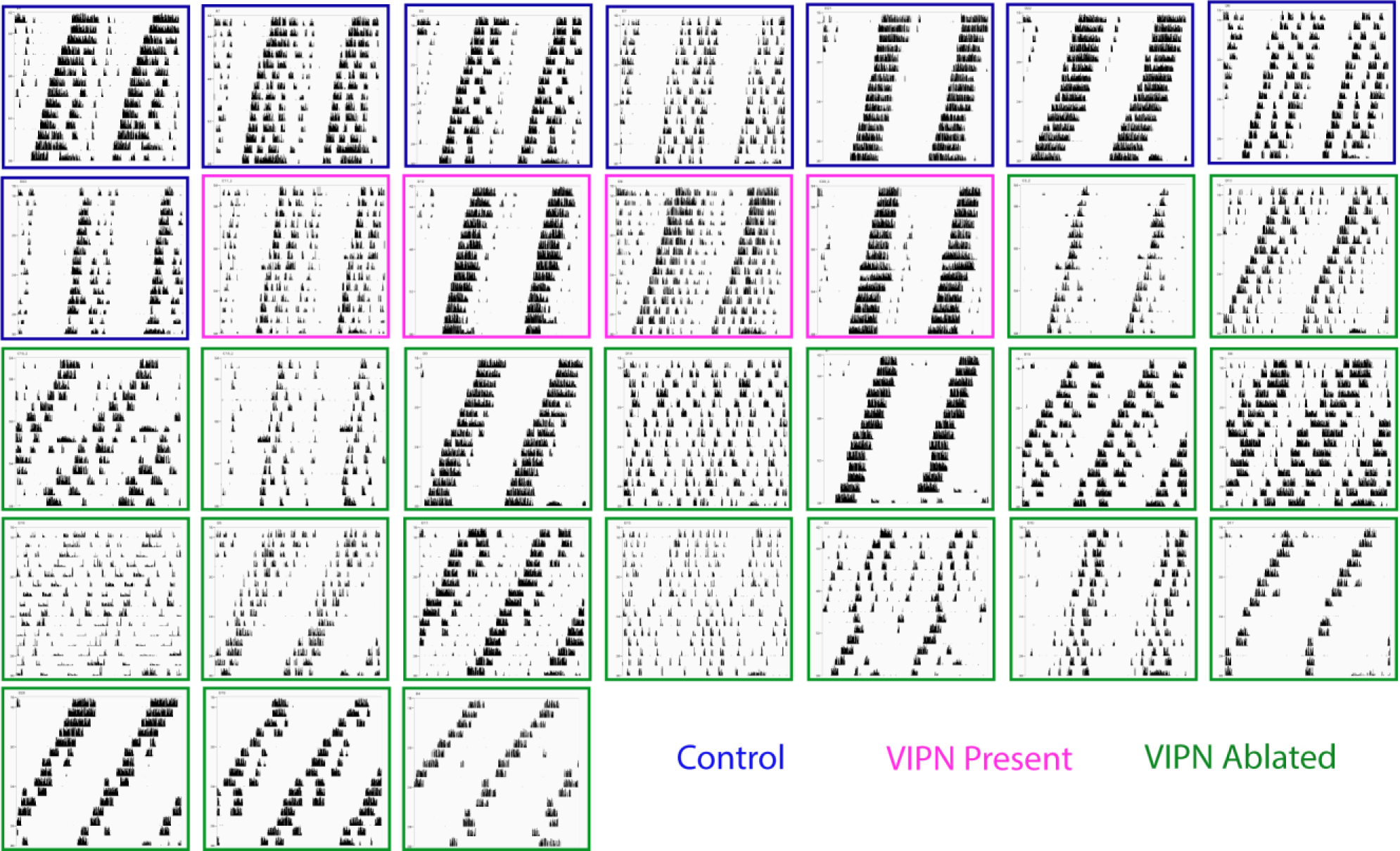
Locomotor Phenotype in constant darkness. Locomotor Actograms during constant darkness ordered by level of VIP staining (top-left to bottom-right). Note that the predominant phenotype among mice with the lowest level of VIP staining was a shortened period and a decreased duration of activity.

**Figure 3.**
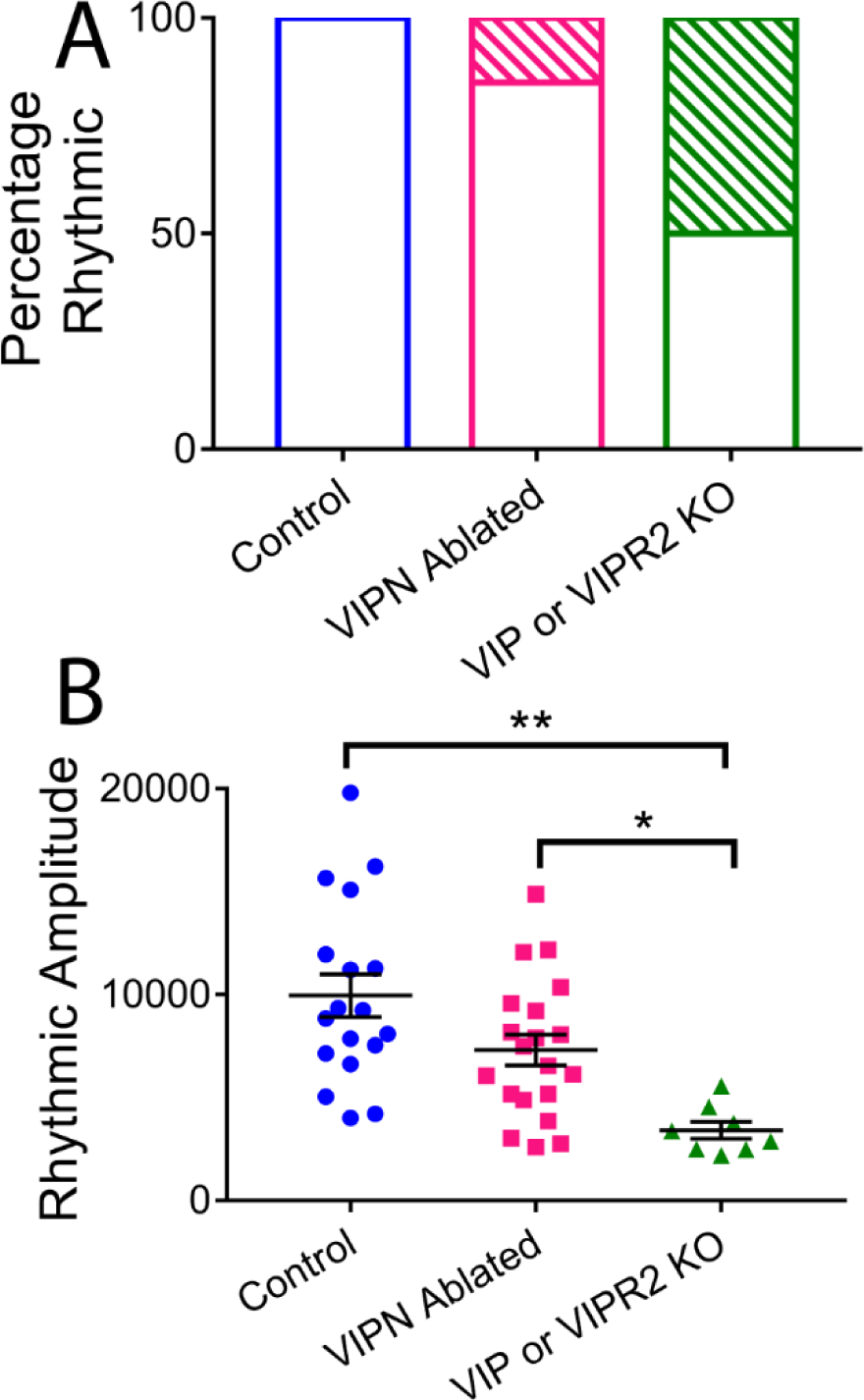
Comparison of VIP SCN neuronal ablation to genetic knockout of *Vip* or *Vipr2*. **A)** Under constant darkness only 15% of VIPN Ablated mice are behaviorally arrhythmic (17/20), compared to 50% of *Vip* or *Vipr2* knockout mice (4/8). **B)** Rhythmic amplitude is substantially decreased in *Vip* or *Vipr2* knockout mice compared to control mice or mice lacking VIP neurons.

### Ablation of VIP neurons dampens the daily surge in corticosterone

Following locomotor activity recordings, a subset of mice was placed in custom-built cages that allow frequent, noninvasive measurement of fecal corticosterone levels. Mice habituated to the cages for 2 days in total darkness before fecal collection every 4 h for two more days (n = 13 VIPN ablated, n = 10 Control). When we aligned individual corticosterone rhythms to the time of their first daily peak, we found that mice lacking VIP neurons had dampened circadian corticosterone levels compared to controls, with roughly half the average (29.77±2.43 vs. 41.86±4.20 ng corticosterone per mg feces, t(21) = 2.63, p = 0.0157), maximum (48.32±3.84 vs. 94.36±9.93 ng corticosterone per mg feces, t(11.69) = 4.32, p = 0.001), and peak-to-trough daily amplitude of corticosterone levels (30.37±3.25 vs. 74.06±9.50 ng/mg, t(11.12) = 4.35, p = 0.0011, Figure 4). We found no difference in the daily minimum of corticosterone (17.04±2.19 vs. 20.18±3.11 ng/mg, t(21) = 0.85, p = 0.4042) and conclude that SCN VIP neurons are necessary for the daily surge in corticosterone.

**Figure 4.**
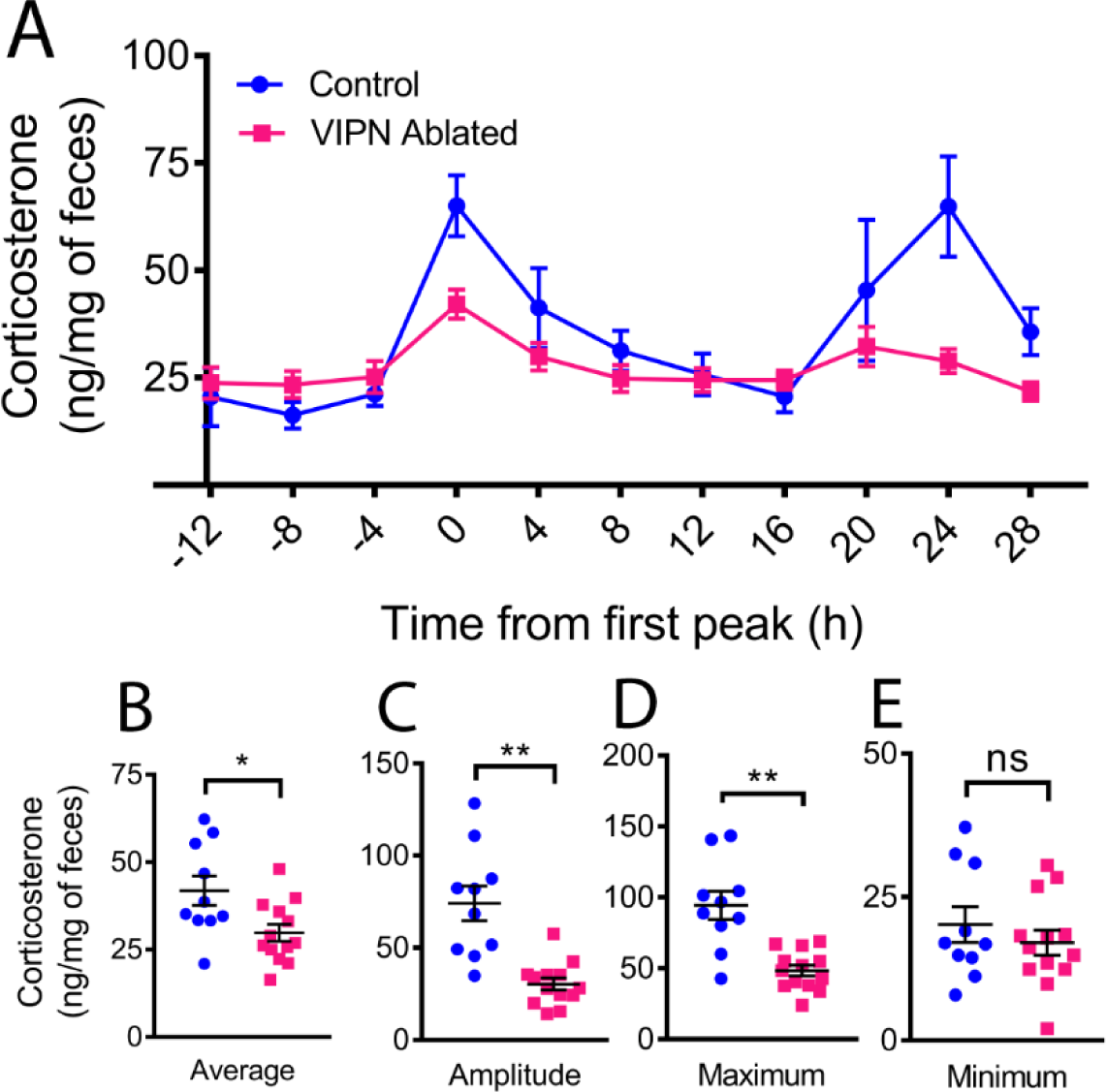
Ablation of VIP neurons dampens the circadian rhythm in corticosterone. **A**) Average profile of fecal corticosterone levels over two days in constant darkness from mice with (blue) and without (magenta) SCN VIP neurons. Individual traces were phase-aligned to their first daily peak. **(B)** Loss of VIP neurons reduced daily average, (**C**) peak-to-trough amplitude of corticosterone and (**D)** maximum, (**E**) but did not change the daily minimum.

### Circadian Phenotype correlates with VIP Staining Intensity

To measure the extent of VIP neuron ablation, we quantified VIP immunofluorescence staining. All mice were sacrificed during entrained conditions (12h:12h light-dark cycle) during late subjective day (CT 7-10) when arginine vasopressin (AVP) staining is high (Jin et al., 1999; Silver et al., 1999; Tominaga, Shinohara, Otori, Fukuhara, & Inouye, 1992). On average, VIP-IRES-Cre mice injected with Cre-dependent Caspase3 had significant reductions in VIP staining intensity compared to control mice (110.9±43.6 compared to 483.4±98.9 relative VIP staining intensity, t(27) = 3.768, p = 0.0008), but no changes in AVP intensity (791.3±8.3 compared to 770.6±57.2 relative AVP staining intensity, t(23.65) = 0.2081, p = 0.8369; Figures 5A-B). Among the mice that received the virus into the SCN, we found a subset (n=4) with VIP staining levels comparable to controls (“VIPN present” mice). Across all mice, staining intensity for VIP in the SCN strongly correlated with increasing period (r(24) = 0.40, p = 0.0005, Figure 5C) and decreasing onset variability (r(24) = 0.3485, p = 0.0015, Figure 5D) of daily locomotor rhythms in constant darkness. The three VIPN ablated mice that were behaviorally arrhythmic had comparable levels of VIP staining to other VIPN ablated mice (see orange stars, Figures 5C-D). AVP staining intensity did not correlate with locomotor period (r(24) = 0.01, p = 0.55) or onset variability (r(24) = 0.09, p = 0.14). We conclude that reducing the number of VIP neurons in the SCN results in reduced circadian period and reduced cycle-to-cycle precision in locomotor rhythms.

**Figure 5.**
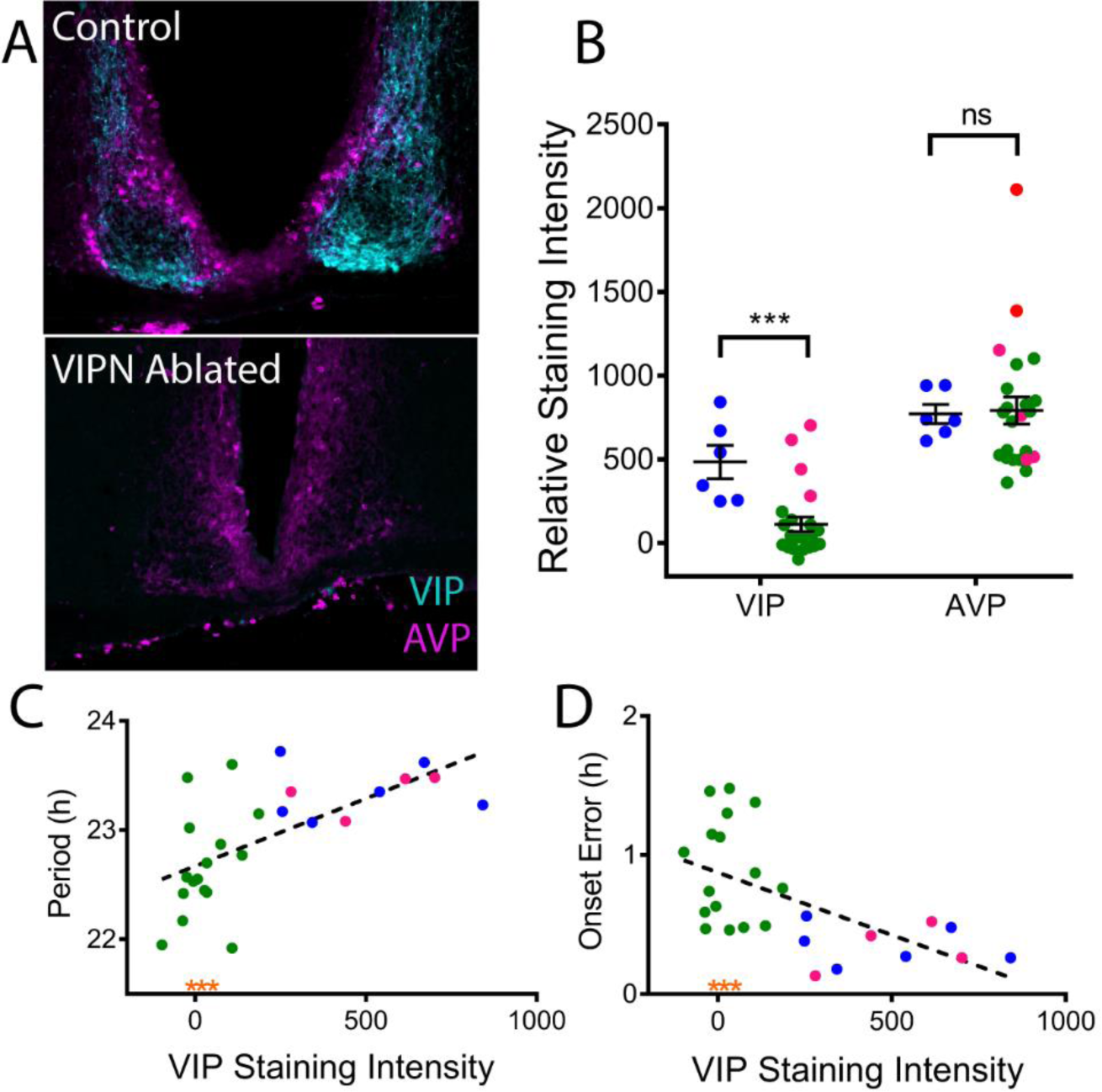
Locomotor Phenotype Correlates with Level of VIP expression. **A)** Representative coronal sections of the SCN from control and VIPN ablated mice double labelled for VIP (cyan) and AVP (magenta). Note the loss of VIP immunoreactivity in the mouse treated with AAV-Casp3. **B)** Quantification of VIP and AVP immunostaining revealed lower average VIP, but not AVP, immunostaining in the SCN of mice treated with AAV8-Casp3 (green: VIPN ablated and magenta: VIPN present) compared to controls (blue dots). **C)** Circadian period length correlated with VIP staining intensity while (**D**) day-to-day onset variability inversely correlated with VIP staining intensity. Note that VIPN ablated mice exhibiting arrhythmic circadian behavior have VIP staining intensities in line with other VIPN ablated mice (orange asterisks)

### In vitro deletion of VIP neurons impairs circadian rhythmicity in the neonatal SCN

To further compare the effects of loss of VIP to loss of VIP neurons, we generated triple transgenic mice that expressed a reporter of the clock protein, PERIOD2 (PER2) in all cells and red fluorescence only in VIP neurons (VIP-IRES-Cre/+, floxed-ROSA-tdTOMATO/+, PERIOD2::Luciferase/+). At postnatal day 6, we dissected the SCN and infected SCN slices with either AAV-DIO-Caspase3 to kill VIP neurons or a control virus (AAV-DIO-eYFP). We then recorded PER2-driven bioluminescence every 10 minutes for up to a month while imaging fluorescence once a week. By three weeks after viral expression, caspase-treated SCN slices showed significantly fewer VIP neurons than control slices (18±5 compared to 268±28, t(5.327) = 8.71, p = 0.0002; Figure 6). In VIPN ablated SCN slices, the average (127,813±7770 compared to 229,559±20,699 counts/10 min, t(13) = 5.8, p < 0.0001), and daily peak-to-trough amplitude of PER2 expression (36,641±8897 compared to 132,085±14,425 counts/10 min, t(13) = 5.794, p < 0.0001) were both dramatically reduced. Notably, the quality of the rhythm also was reduced (0.91±0.03 compared to 0.99±0.00, t(7.026) = 2.46, p = 0.0434, as assessed by the correlation coefficient of a cosine fit to each bioluminescence trace), although the circadian period was not (24.04±0.21 h compared to 23.64±0.17h, t(13) = 1.42, p = 0.1776). We conclude that loss of VIP neurons shortly after birth reduces the amplitude and precision of circadian rhythms in the SCN. These results are consistent with the reduced synchrony of circadian cells in the SCN of *Vip* and *Vipr2* null mice.

**Figure 6.**
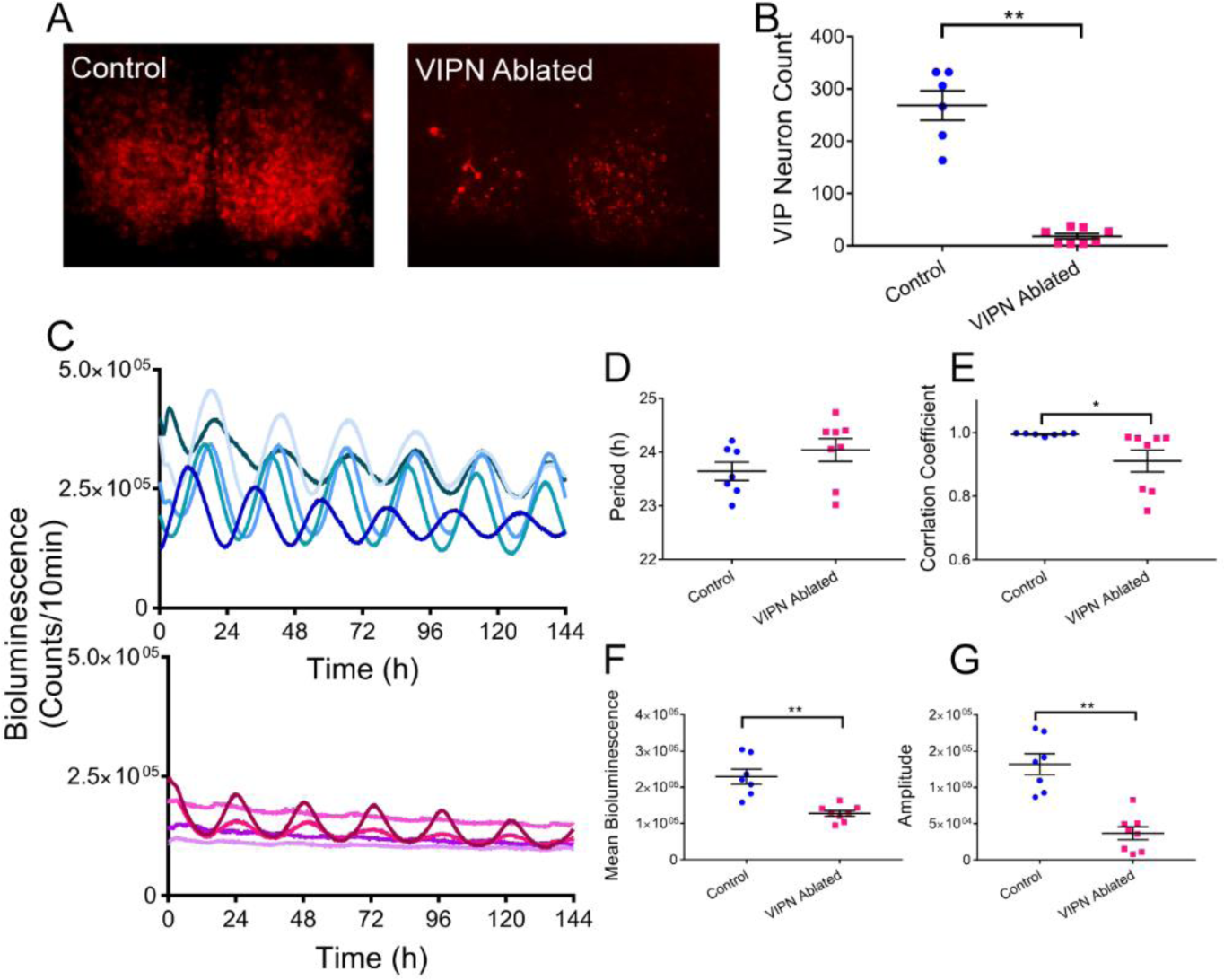
Ablation of VIP Neurons in neonatal SCN slices *in vitro* decreases circadian rhythmicity. **A)** VIP-specific tdTOMATO fluorescence was substantially decreased in SCN slices treated with AAV-DIO-Caspase3 (VIPN ablated) compared to those treated with an eYFP-expressing virus (control). **B)** Quantification of the number of VIP neurons remaining in the SCN slices 3 weeks after treatment. **C)** PER2 expression over days from individual SCN slices 3 weeks after treatment with control (top blue traces) or caspase3 (bottom magenta traces) viruses. **D)** SCN slices lacking VIP neurons showed no change in circadian period, **E)** but reduction in rhythm quality as measured by the correlation coefficient of a cosine fit, **F)** mean bioluminescence, and **G)** peak-to-tough circadian amplitude.

## Discussion

We found that ablation of VIP SCN neurons in adult mice caused a circadian phenotype distinct from that seen in *Vip-* or *Vipr2-* null mice. Whereas adult loss of VIP neurons has relatively little effect on circadian rhythms in locomotion compared to the arrhythmicity seen in mice deficient for VIP signaling from birth, both manipulations result in a blunting of glucocorticoid rhythms. Combined with the striking decreases in circadian gene expression and rhythmicity seen when VIP neurons are ablated in the neonatal SCN, we conclude that VIP SCN neurons may have differential roles in development and adulthood.

The absence of a severe circadian locomotor phenotype in VIPN-ablated, adult mice may suggest that VIP signaling plays a distinct role in the development of the circadian system. VIP neuropeptide expression within the SCN is first detected in early postnatal days followed by retinal innervation of the SCN (Carmona-Alcocer, Abel, et al., 2018)(Carmona-Alcocer, Rohr, Joye, & Evans, 2018; Sernagor, 2005). VIP may, thus, be necessary for the correct formation of retinal input to the SCN, intra-SCN connectivity and SCN output to hypothalamic areas. It is possible that disruption of VIP signaling during this critical postnatal period contributes to the severe circadian disruption seen in *Vip* or *Vipr2* deficient mice. This appears different from findings in *Drosophila*, where deletion of the neuropeptide PDF gene or cells produces an identical circadian behavioral phenotype. In mammals, VIP neuropeptide signaling may play a role in establishing circadian circuit features during development as it does in modulating the excitatory/inhibitory balance in visual cortex (Batista-Brito et al., 2017).

Because we found that deletion of VIP neurons in the neonatal SCN results in similar reductions in circadian rhythms in the SCN to previously published genetic or pharmacological abrogation of VIP signaling (Aton et al., 2005; T. M. Brown et al., 2007; Colwell et al., 2003; Harmar AJ et al., 2002; Maywood et al., 2006), we conclude that these neurons are important for synchrony among circadian cells early in development, but not in the adult. This is consistent with evidence that other signals, including the neuropeptides AVP or Gastrin Releasing Peptide, can act to synchronize circadian cells in the SCN (Timothy M. Brown et al., 2005; Herzog, Hermanstyne, Smyllie, & Hastings, 2017; Maywood et al., 2011). Furthermore, mice that lose VIP neurons in adulthood share features with the subset of mice born deficient for Vip signaling that retain circadian rhythms such as significantly shortened periods. This indicates that SCN VIP neurons normally serve to lengthen the circadian period, consistent with results showing that chronic stimulation of VIP receptors lengthens circadian period in vivo (Pantazopoulos, Dolatshad, & Davis, 2010), VIP largely phase delays the *in vitro* SCN (An, Irwin, Allen, Tsai, & Herzog, 2011; Reed, Meyer-Spasche, Cutler, Coen, & Piggins, 2001), activation of SCN VIP neurons can phase delay behavior *in vivo* (Mazuski et al., 2018), and that the phase delaying effects of light require activation of SCN VIP neurons (Jones, Simon, Lones, & Herzog, 2018). Our results also suggest that VIP neurons are necessary for circadian locomotor activity in the second half of the circadian night. Ablation of VIP neurons compresses circadian locomotor activity to the first part of the subjective night.

We found reduced corticosterone levels in mice lacking SCN VIP neurons, suggesting that they underlie stimulation of the daily surge in glucocorticoids. Prior results showing loss of glucocorticoid rhythms in VIP null mice (Loh et al., 2008) can now be interpreted as likely caused by disruption of VIP SCN neurons and not VIP in other areas (such as the adrenal glands) or arrhythmic circadian behavior. Neuronal projections from the SCN to the paraventricular nucleus of the hypothalamus (PVN) are necessary for the daily rhythm in glucocorticoids (Buijs, Kalsbeek, van der Woude, van Heerikhuize, & Shinn, 1993), and SCN VIP neurons project to the PVN (Abrahamson & Moore, 2001). It remains to be tested whether activity in these VIP projections is sufficient and necessary for daily regulation of glucocorticoid release. Future work should further elucidate the relative roles of VIP in establishing and maintaining the many daily rhythms in physiology and behavior.

Light-Dark Analyses (LD)
| Group | Genotype | N | Injection | Period<sup>Y</sup> | Onset Error<sup>Y</sup> | Duration of Activity<sup>X</sup> | Activity Per Day<sup>Y</sup> |
| --- | --- | --- | --- | --- | --- | --- | --- |
| Control<sup>#</sup> | C57Bl/6JN, P2L/+ or VIP-Cre | 11 | AAV-DIO-Caspase3 or ACSF | 24.01 ± 0.02 | 0.25 ± 0.6 | 9.19 ± 0.64 | 19264 ± 2339 |
| VIPN Present | VIP-Cre | 7 | AAV-DIO-Caspase3 | 24.00 ± 0.01 | 0.06 ± 0.02 | 9.46 ± 1.07 | 15612 ± 3185 |
| VIPN Ablated | VIP-Cre | 20 | AAV-DIO-Caspase3 | 24.01 ± 0.02 | 0.29 ± 0.05 | 6.96 ± 0.33* | 16433 ± 1559 |
| Genetic knockout | VIP or VIPR KO | 8<sup>\$1</sup> | N/A | 23.94 ± 0.01* | 0.75 ± 0.25 | 8.21 ± 0.69 | 10113 ± 1958* |

Constant Darkness Analyses (DD)
| Group | Genotype | N | Injection | Period<sup>X</sup> | Onset Error<sup>Y</sup> | Duration of Activity<sup>X</sup> | Activity Per Day<sup>X</sup> |
| --- | --- | --- | --- | --- | --- | --- | --- |
| Control<sup>#</sup> | C57Bl/6JN, P2L/+ or VIP-Cre | 11 | AAV-DIO-Caspase3 or ACSF | 23.42 ± 0.08 | 0.44 ± 0.06 | 10.16 ± 0.52 | 20835 ± 2837 |
| VIPN Present | VIP-Cre | 7 | AAV-DIO-Caspase3 | 23.47 ± 0.09 | 0.49 ± 0.10 | 8.92 ± 0.80 | 19320 ± 2942 |
| VIPN Ablated | VIP-Cre | 20<sup>\$2</sup> | AAV-DIO-Caspase3 | 22.69 ± 0.12** | 0.87 ± 0.09* | 7.17 ± 0.54** | 14290 ± 1861 |
| Genetic knockout | VIP or VIPR KO | 7<sup>\$3</sup> | N/A | 23.15 ± 0.18 | 1.85 ± 0.40** | 9.83 ± 0.70 | 15929 ± 3923 |

<sup>#</sup> No statistical differences between non-cre expressing control mice injected with AAV8-DIO-Caspase3 (n = 6) or VIP-Cre control mice injected with ACSF (n = 5)
<sup>\$1</sup> N = 7 for period due to arrhythmicity of 1 mouse
<sup>\$2</sup> N = 17 for period, onset error and duration of activity analyses due to arrhythmicity in 3 mice
<sup>\$3</sup> N = 4 for period, onset error and duration of activity due to arrhythmicity in 4 mice
<sup>X</sup> Ordinary one-way ANOVA with multiple comparisons to control group
<sup>Y</sup> Non-parametric Kruskal-Wallis test
\* p < 0.05
\*\* p < 0.01

## Acknowledgements

This work was supported by NIH grants UO1 EB02195601 (E.D.H) and F31-GM11517 (C.M.). The authors thank members of the Herzog lab, especially Drs. Matt Tso and Daniel Granados-Fuentes for technical contributions.

